# New shallow water species of Caribbean *Ircinia* Nardo, 1833 (Porifera: Irciniidae)

**DOI:** 10.1101/2020.09.01.277210

**Authors:** Joseph B. Kelly, Robert W. Thacker

## Abstract

Seven *Ircinia* growth forms were collected from three sites in the Caribbean (Bocas del Toro, Panama; the Mesoamerican Barrier Reef, Belize; and the Florida Keys, United States of America). Previous research used an integrative taxonomic framework to delimit species boundaries among these growth forms. Here, we present descriptions for these species, six of which are new to science (*Ircinia lowi* **sp. nov**., *Ircinia bocatorensis* **sp. nov**., *Ircinia radix* **sp. nov**., *Ircinia laeviconulosa* **sp. nov**., *Ircinia vansoesti* **sp. nov**., *Ircinia rutzleri* **sp. nov**.) in addition to one species *conferre* (*Ircinia* cf. *reteplana* Topsent, 1923).

## Introduction

*Ircinia* Nardo, 1833 is a genus of sponges diagnosable from other genera in the family Irciniidae Gray, 1867 by the possession of cored fascicular primary fibers and the lack of cortical armoring (Hooper & van Soest, 2002). The study of *Ircinia* taxonomy has been historically problematic as *Ircinia*, like other members of the order Dictyoceratida Minichin, 1900, are aspiculate and present few anatomical features that can be used to infer relatedness and taxonomic boundaries among species (Erpenbeck et al., 2020). *Ircinia* also display a considerable degree of morphological plasticity that confounds the identification of features that are representative of a given species (de C. Cook & Bergquist, 1999). Single locus genetic barcoding has likewise seen limited success, as the loci that are typically used to provide species- and population-level phylogenetic resolution in metazoans, which include the *cytochrome oxidase c subunit 1* (CO1) and the internal transcribed spacer set (ITS), typically are either incompletely sorted or largely invariant among nominal species of *Ircinia* (Pöppe et al., 2010; Riesgo et al., 2016; Kelly & Thacker 2020, *in review*). The pitfalls that arise from the use of morphological data alone or by restricting genetic data to a single locus or a few loci, which is also a questionable practice on the basis of the high probability of gene tree and species tree discordance (Degnan & Rosenberg, 2006), necessitate the implementation of integrative taxonomic research frameworks within *Ircinia* that include genome-wide evidence of species boundaries.

Shallow water environments of the Caribbean are commonly inhabited by three nominal species of *Ircinia*: *I. campana*; *I. strobilina* Lamarck, 1816; and *I. felix* Duchassaing & Michelotti, 1864. Alongside these three species can be found several *Ircinia* growth forms that are recognizable in the field based on the overall shape of their bodies, the spacing and height of their conules, and the position and size of their oscula (Diaz, 2005; Erwin & Thacker, 2007; Rützler et al., 2000; van Soest, 1978; Wulff, 1994, 2013). The growth forms are also generally regarded as being ecologically distinct in that they contain different tissue densities of chlorophyll *a* (Erwin & Thacker, 2007), can exhibit habitat preference, and have distinct microbiome compositions (Kelly & Thacker 2020, *in review*; Kelly et al. 2020, *in review*). Recently, genetic species boundaries were delimited among several of these growth forms using Bayesian species delimitation (BFD*) with genome-wide SNP data (Leaché et al., 2014, Kelly & Thacker 2020, *in review*; Kelly et al. 2020, *in review*). On the basis that these growth forms are genetically and morphologically distinct and appear ecologically divergent as evidenced by their possession of unique microbiomes, we designate the growth forms as new species to science and provide taxonomic descriptions.

## Methods

*Ircinia* specimens were collected from three sites in the Caribbean: four growth forms were collected from Bocas del Toro, Panama; three from the Mesoamerican Barrier Reef, Belize; and one from the Florida Keys, United States of America (Figure 1, Table 1). Each of the growth forms are specific to a given habitat type and are found in either coral reefs, seagrass beds, or on mangrove prop roots. Tissue samples were fixed in 4% paraformaldehyde (PFA) that was prepared by diluting 32% PFA stock in filtered seawater. The 4% PFA solution was replaced at the 24- and 48-hour marks to ensure complete irrigation of the tissue. Histological sections were inspected using compound light microscopy.

**Table 1.**
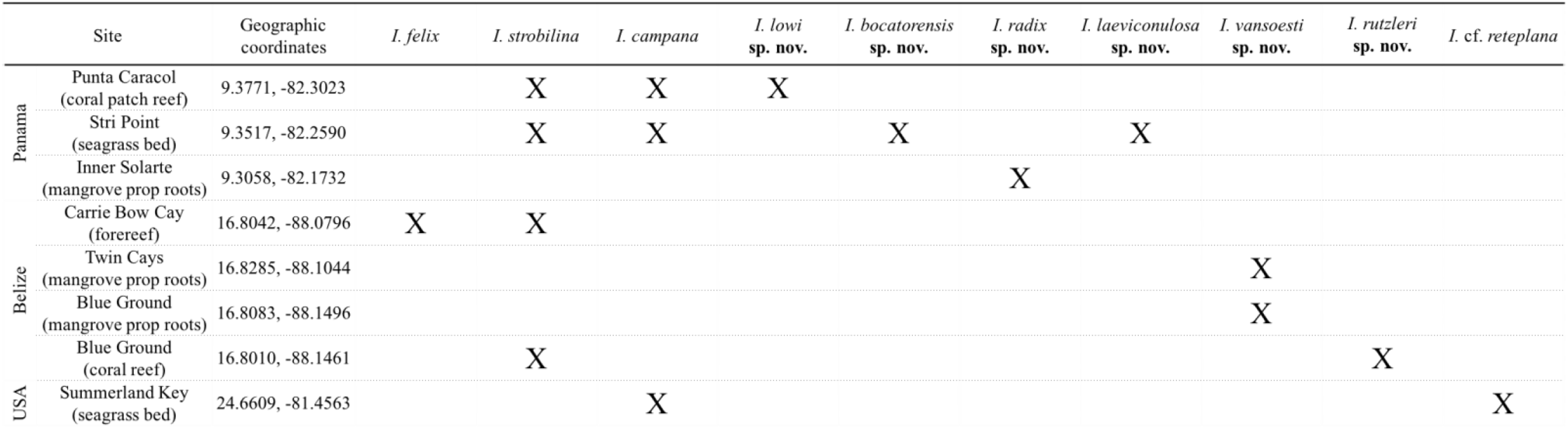
Sampling locations of *Ircinia* spp.

**Figure 1.**
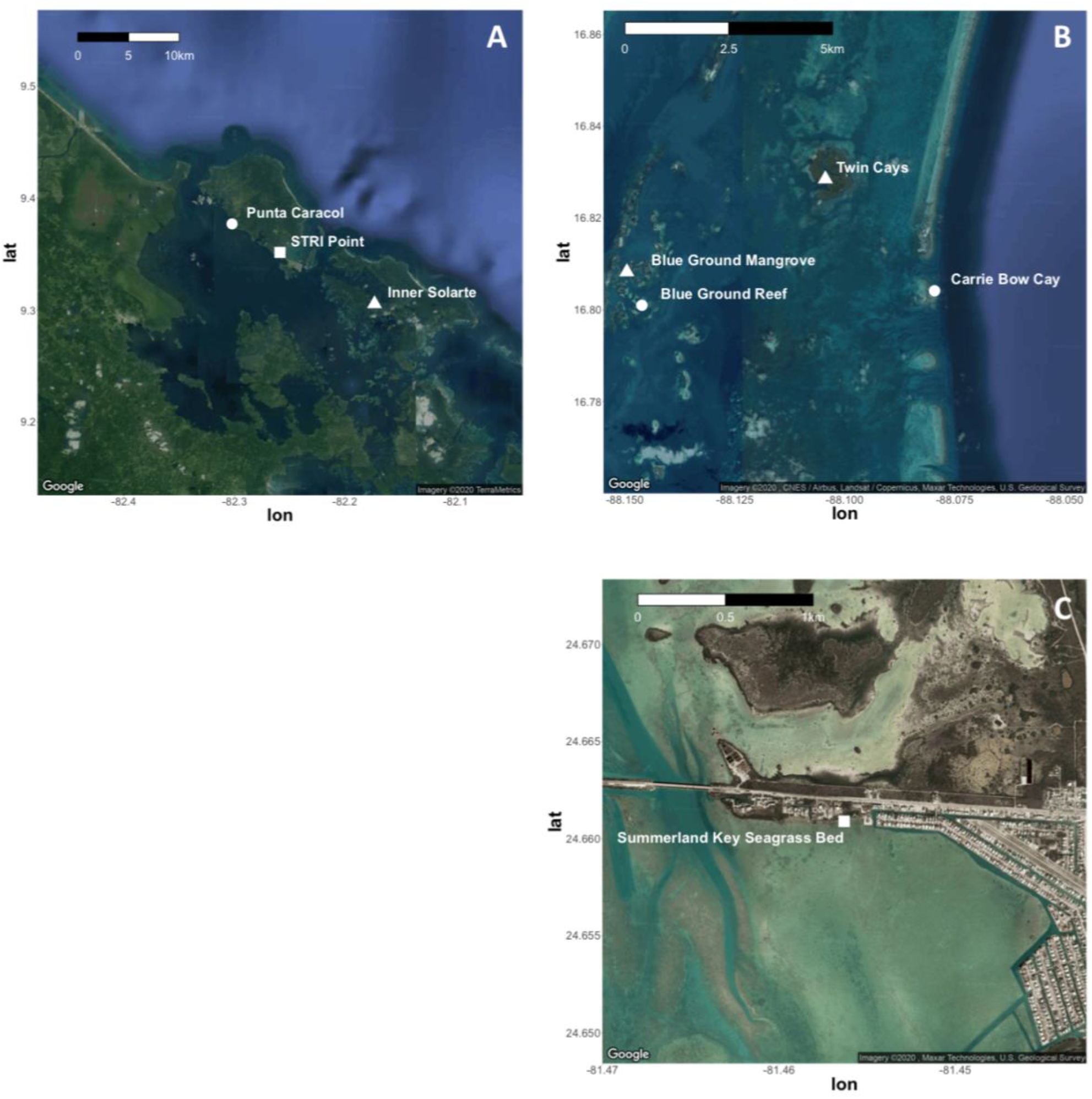
Maps of sampling locations. **A:** Bocas del Toro, Panama; **B:** Mesoamerican Barrier Reef, Belize; **C:** Summerland Key, United States of America. Circles are coral reefs or coral patch reefs, squares are seagrass beds, and triangles are mangroves.

## Results

### General remarks

The characters that distinguish *Ircinia* from other irciniid genera hold for the *Ircinia* spp. nov., in that the current *Ircinia* have cored fascicular primary fibers and lack cortical armoring. However, we note that the cortices of the *Ircinia* spp. nov. sometimes incorporate foreign spicules and sand. Foreign spicules, when present in the mesohyl, fibers, or cortex, are predominantly oxea and style fragments.

The dimensions of the skeletal fibers and conule heights of the *Ircinia* spp. nov. below are dissimilar from *I. campana* (Lamarck, 1814), *I. strobilina* (Lamarck, 1816), and *I. felix* (Duchassaing & Michelotti, 1864) (Table 2). Additionally, all lack the characteristic dermal reticulation of *I. felix*; their conules instead arise from smooth dermal surfaces. The body shape of *I. reteplana* is distinct from those of the aforementioned *Ircinia* in that it is composed of flattened, interconnecting branches, with the exception of the Floridian growth form. Because the Floridian *Ircinia* growth form (called ‘Ramose’ in Kelly & Thacker 2020, *in review* and Kelly et al. 2020, *in review*) often displays a flattened branching morphology, we designate this growth form as *Ircinia* cf. *reteplana* Topsent, 1923. However, *Ircinia* cf. *reteplana* can also possess a rounded branching morphology and the branches of *Ircinia* cf. *reteplana* seldom interconnect. This growth form represents either a new species of *Ircinia* or it represents an extension of the documented range of *I. reteplana* to the Florida Keys. For reference, the range of *I. reteplana* encompasses the entirety of the Antilles and spans the Caribbean to the coast of Venezuela, and extends to the tip of the Yucutan Peninsula (Van Soest et al., 2019). The collections of *I. strobilina, I. felix*, and *I. campana* (Kelly & Thacker 2020, *in review*; Kelly et al. 2020, *in review*) were made within the documented ranges of these species (Table 1).

**Table 2.** A morphological comparison of shallow water Caribbean *Ircinia* (van Soest, 1978).

| Species | Massive fascicular fiber width (μm) | Inter-connecting fiber width (μm) | Irciniid fiber width (μm) | Conule height (mm) | Oscula diameter (cm) |
| --- | --- | --- | --- | --- | --- |
| *Ircinia campana*<br>Lamark, 1816 | 300-700 | 30-150 | 3-6 | ~ 4 | 0.4-1 |
| *Ircinia strobilina*<br>Lamark, 1816 | 200-300 | 20-60 | 4-6 | 2-15 | 0.4-1 |
| *Ircinia felix*<br>Duchassaing & Michelotti, 1864 | 200-550 | 15-100 | 2-6 | 1-2 | 0.1-0.8 |
| *Ircinia reteplana*<br>Topsent, 1923 | not reported | 25-100 | ~6 | not reported | 0.5-0.8 |
| *Ircinia lowi*<br>**sp. nov.** | 80-200 | 20-60 | 6-11 | > 2 | > 0.5 |
| *Ircinia bocatorensis*<br>**sp. nov.** | 200-520 | 15-65 | 6-10 | 3-5 | > 1.5 |
| *Ircinia laeviconulosa*<br>**sp. nov.** | 110-160 | 20-50 | 4-6 | 1.5-1.75 | > 1.2 |
| *Ircinia radix*<br>**sp. nov.** | 110-250 | 10-50 | 6-8 | ~2 | > 1.2 |
| *Ircinia vansoesti*<br>**sp. nov.** | 90-300 | 25-60 | 7.5-9 | 1-1.5 | 0.5-1.2 |
| *Ircinia rutzleri*<br>**sp. nov.** | 120-240 | 30-50 | 7.5-9 | 1-1.3 | 0.7-1 |
| *Ircinia* cf. *reteplana*<br>Topsent, 1923 | 90-200 | 20-80 | 7.5-9 | 1-1.2 | ~ 0.5 |

### Systematics

**Phylum Porifera Grant, 1836**

**Class Demospongiae Sollas, 1885**

**Subclass Keratosa Grant, 1861**

**Order Dictyoceratida Minchin, 1900**

**Family Irciniidae Gray, 1867**

**Genus *Ircinia* Nardo, 1833**

## *Ircinia lowi* sp. nov

**Holotype: P16×42 (USNM 1582268)**.

**Paratypes: P16×41 (USNM 1582267), P16×43 (USNM 1582269), P16×44 (USNM 1582270), P16×52 (USNM 1582278)**.

**Type locality: Bocas del Toro, Panama**

### Diagnosis

*Ircinia* that form a thickly encrusting growth habit, with a forest green external surface color. Sometimes possesses digitate projections of the body (Figure 2).

**Figure 2.**
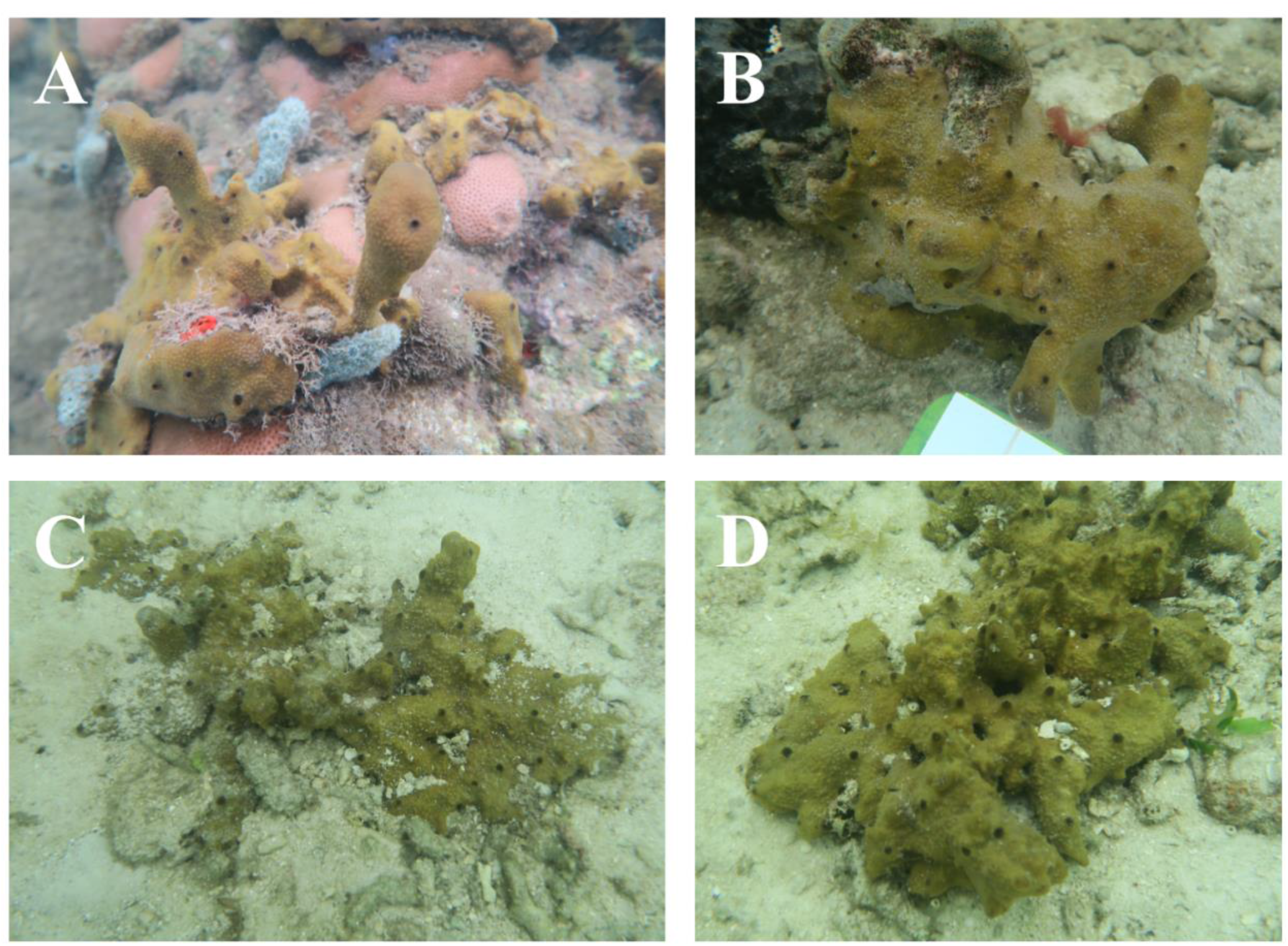
*Ircinia lowi* sp. nov. **A:** USNM 1582268 (holotype), **B:** USNM 1582267 (paratype), **C:** USNM 1582269 (paratype), **D:** USNM 1582270 (paratype).

### External morphology

Conules small (<2mm in height), sometimes of lighter color than the rest of the body, similar to *I. laeviconulosa* sp. nov. and *I. radix* sp. nov., although slightly sharper. The body is dotted with black oscula that are uniform in size and either sit at a slight relief or are flush, usually 0.5 cm or less in diameter.

### Interior morphology

Massive fascicular fibers 80-200 μm wide, cored. Interconnecting fibers 20-60 um wide, sparsely cored. Irciniid filaments 6-11 μm wide.

### Ecology

All specimens were collected from shallow depths (0.4 – 0.5 m) on patch reefs that occur in association with small seagrass beds.

### Etymology

This species is named for the immunologist Jun Siong Low.

### Remarks

Tissue takes on a slightly crisper consistency when preserved in ethanol for several days. Referred to as the ‘Encrusting’ growth form in Kelly & Thacker (2020) and Kelly *et al*. (2020).

## *Ircinia bocatorensis* sp. nov

**Holotype: P16×58 (USNM 1582284)**.

**Paratypes: P16×56 (USNM 1582282), P16×61 (USNM 1582287), P16×63 (USNM 1582289), P16×65 (USNM 1582291), P16×69 (USNM 1582295)**.

**Type locality: Bocas del Toro, Panama**

### Diagnosis

*Ircinia* with a massive, sometime cone-like growth morphology and a tan exterior color (Figure 3).

**Figure 3.**
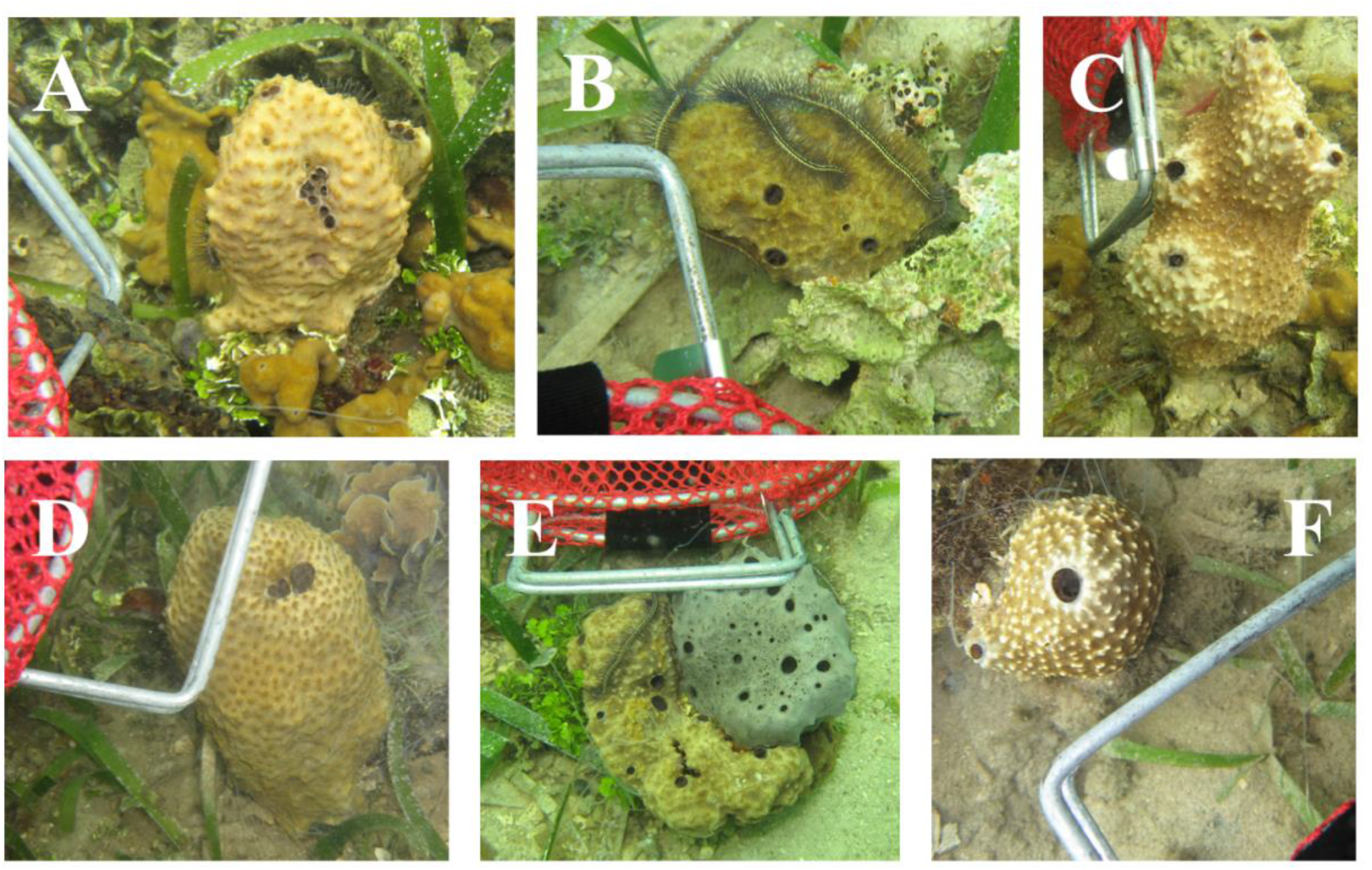
*Ircinia bocatorensis* sp. nov. **A:** USNM 1582284 (holotype), **B:** USNM 1582289 (paratype), **C:** USNM 1582287 (paratype), **D:** USNM 1582295 (paratype), **E:** USNM 1582291 (paratype) adjacent to *I. strobilina*, **F:** USNM 1582282 (paratype).

### Exterior morphology

Conules 3-5 mm in height, dully sharp to knobby, typically darker or lighter in color than the rest of the sponge body. Oscula usually black, either flush or positioned at the ends of cone-like outgrowths, of varying diameters but seldom wider than 1.5 cm.

### Internal morphology

Massive fascicular fibers 200-520 μm wide, sparsely cored. Interconnecting fibers 15-65 μm wide, uncored. Irciniid filaments 6-10 μm wide.

### Ecology

All specimens were collected from 0.4 – 1 m depth on a *Thalassia* bed interspersed with small coral colonies.

### Etymology

The species is named for the Panamanian province Bocas del Toro.

### Remarks

One specimen was observed growing in physical contact with *I. strobilina*.

Referred to as the ‘Massive B’ growth form in Kelly & Thacker (2020) and Kelly *et al*. (2020).

## *Ircinia radix* sp. nov

**Holotype: P16×32 (USNM 1582258)**.

**Paratypes: P16×31 (USNM 1582257), P16×33 (USNM 1582259), P16×34 (USNM 1582260)**.

**Type locality: Bocas del Toro, Panama**

### Diagnosis

*Ircinia* with a massive growth form and light pink pinacoderm (Figure 4).

**Figure 4.**
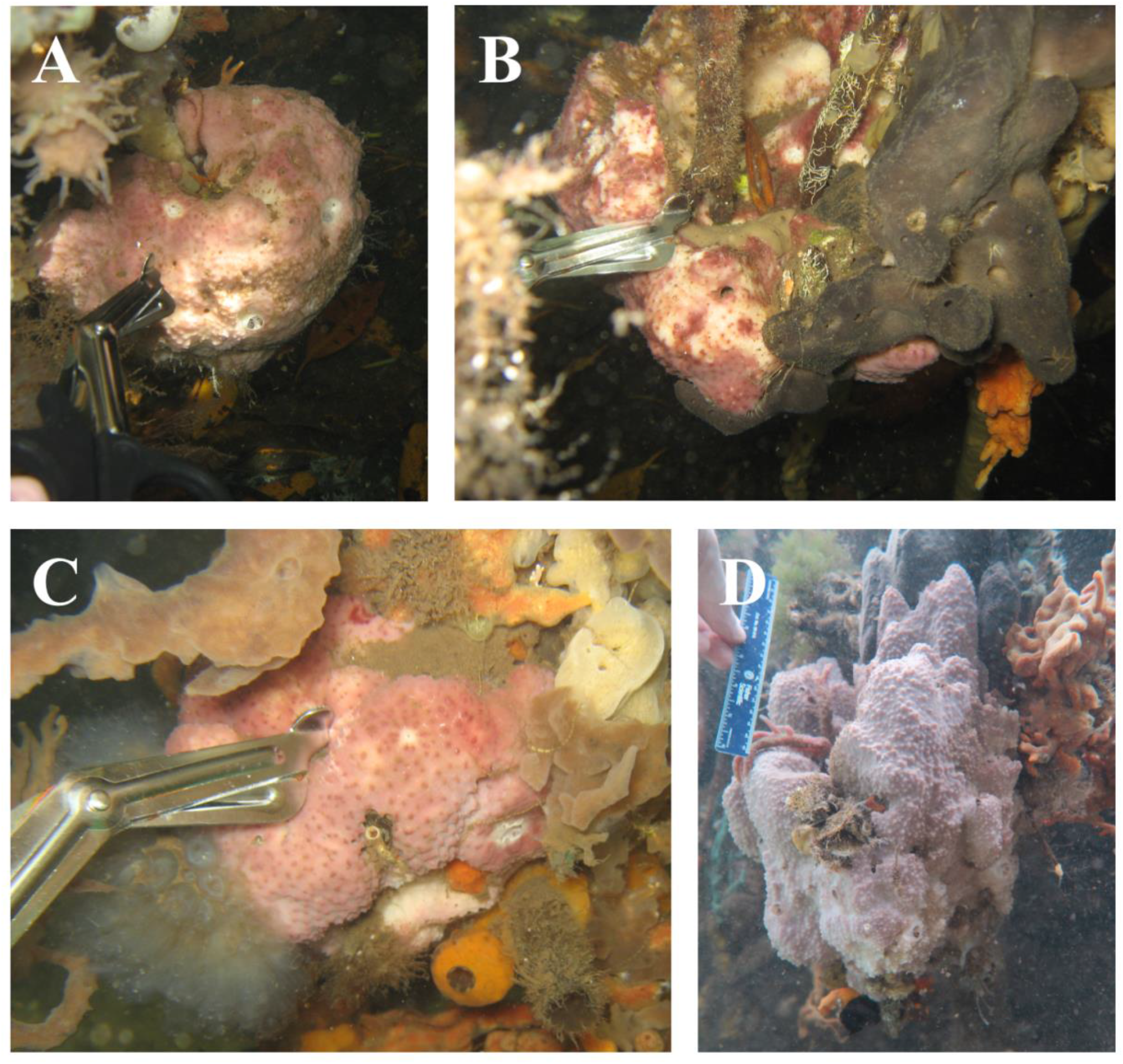
*Ircinia radix* sp. nov. **A:** USNM 1582258 (holotype), **B:** USNM 1582259 (paratype), **C:** USNM 1582260 (paratype), **D:** USNM 1582257 (paratype).

### External morphology

Surface is relatively smooth with low, rounded conules (2 mm). Oscula flush or slightly recessed, occasionally lighter than the exterior of the sponge, usually no larger than 1.2 cm in diameter.

### Interior morphology

Massive fascicular fibers 110-250 μm wide, cored. Interconnecting fibers 10-50 um wide, sparsely cored. Irciniid filaments 6-8 μm wide. Mesohyl and pinacoderm contain haphazardly scattered spicule fragments, with concentrations of foreign inclusions higher in the pinacoderm.

### Ecology

This species inhabits shaded entanglements of mangrove roots. Specimens were collected from depths of 0.5-0.75 m.

### Etymology

The name refers to the mangrove roots that this species lives on.

### Remarks

Growth morphology can range from a round ball (p16×32-34) to an elongated massive form (p16×31). Referred to as the ‘Massive A pink’ growth form in Kelly & Thacker (2020) and Kelly *et al*. (2020).

## *Ircinia laeviconulosa* sp. nov

**Holotype: P16×57 (USNM 1582283)**.

**Paratypes: P16×59 (USNM 1582285), P16×60 (USNM 1582286), P16×62 (USNM 1582288)**.

**Type locality: Bocas del Toro, Panama**

### Diagnosis

*Ircinia* with a massive growth form and dark green pinacoderm (Figure 5).

**Figure 5.**
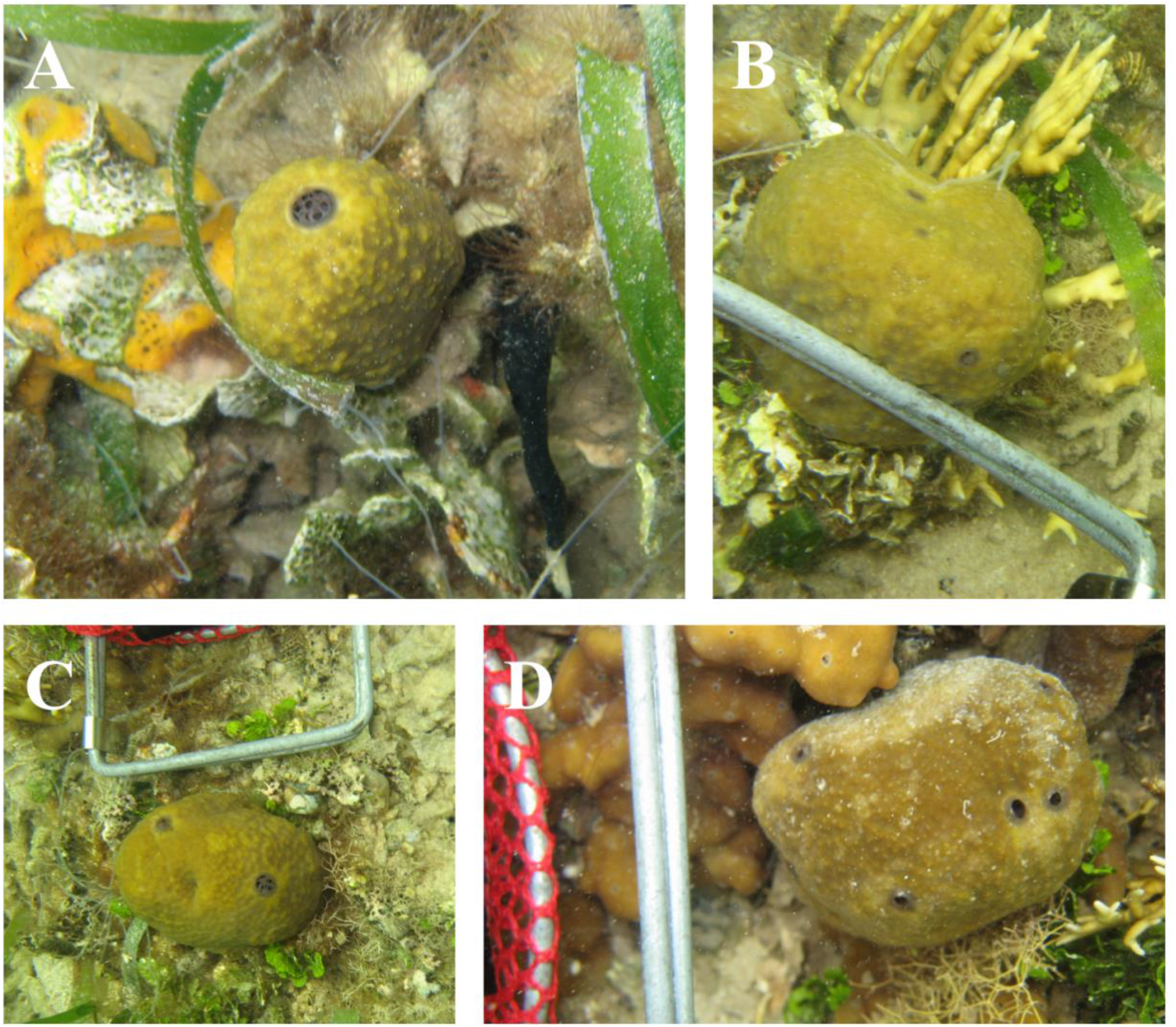
*Ircinia laeviconulosa* sp. nov. **A:** USNM 1582283 (holoype), **B:** USNM 1582285 (paratype), **C:** USNM 1582288 (paratype), **D:** USNM 1582286 (paratype).

### External morphology

Surface texture is similar to *I. radix* sp. nov., although is smoother with lower conules (1.5-1.75 mm). Oscula flush, sometimes slightly darker than the exterior of the sponge, usually no larger than 1.2 cm in diameter.

### Interior morphology

Massive fascicular fibers 110-160 μm wide, cored. Interconnecting fibers 20-50 um wide, sparsely cored. Irciniid filaments 4-6 μm wide. Mesohyl and pinacoderm contain inclusions that resemble those of *I*.*radix* sp. nov.

### Ecology

This species is found among *Thalassia* spp. and coral patches in shallow depths. Specimens were collected from depths of 1-1.5 m.

### Etymology

The name refers to its smooth surface.

### Remarks

All specimens collected had a globose growth morphology. Referred to as the ‘Massive A green’ growth form in Kelly & Thacker (2020) and Kelly *et al*. (2020).

## *Ircinia vansoesti* sp. nov

**Holotype: JK18×20 (**USNM ######## pending)

**Paratypes: JK18×18 (**USNM ######## pending), **JK18×28 (**USNM ######## pending), **JK18×34 (**USNM ######## pending), **JK18×35 (**USNM ######## pending).

**Type locality: Mesoamerican Barrier Reef, Belize**

**Note: lodging of museum vouchers pending**.

### Diagnosis

*Ircinia* with a massive growth form, occasionally lobate, with pinkish brown or gray pinacoderm (Figure 6).

**Figure 6.**
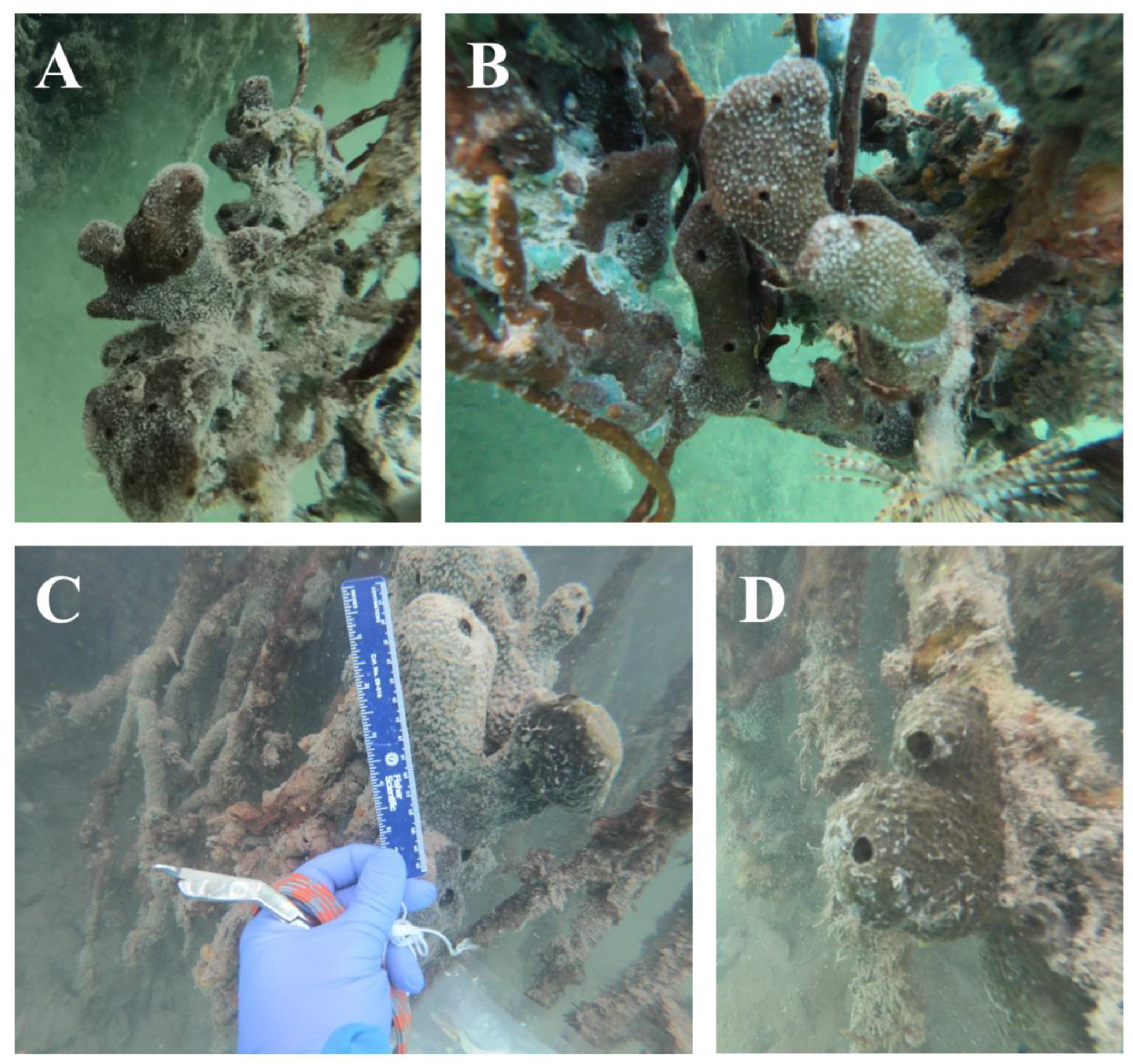
*Ircinia vansoesti* sp. nov. **A:** JK18×20 (holotype), **B**: JK18×18 (paratype), **C:** JK18×28 (paratype), **D**: JK18×35 (paratype).

### External morphology

Surface texture is similar to *I. radix* sp. nov., although with smaller conules (1-1.5 mm). Oscula typically 0.5-1.2 cm in diameter.

### Interior morphology

Massive fascicular fibers 90-300 μm wide, sometimes cored, and always more heavily than interconnecting fibers. Interconnecting fibers 25-60 um wide, usually uncored. Irciniid filaments 7.5-9 μm wide. Cortex usually uncored, sand and foreign spicules occasionally included in mesohyl. Fascicles can be difficult to discern from interconnecting fibers and can be rare.

### Ecology

This species is found growing on *Rhizophora* prop roots at depths of 0.2-1.5 m.

### Etymology

This species is named for the sponge researcher Rob van Soest.

### Remarks

Interior morphology can vary somewhat depending on population, as the Twin Cays specimens contained less foreign inclusions relative to the Blue Ground specimens. This species is also polymorphic with regard to pinacoderm coloration, and multiple color morphs (gray, dark red, dark green) can be found within a population. Referred to as the ‘Sp. 1’ growth form in Kelly *et al*. (2020).

## *Ircinia rutzleri* sp. nov

**Holotype: JK18×23 (**USNM ######## pending)

**Paratypes: JK18×21 (**USNM ######## pending), **JK18×22 (**USNM ######## pending), **JK18×25 (**USNM ######## pending), **JK18×26 (**USNM ######## pending)

**Type locality: Mesoamerican Barrier Reef, Belize**

**Note: lodging of museum vouchers pending.**

### Diagnosis

*Ircinia* with an encrusting growth form and dark gray pinacoderm (Figure 7).

**Figure 7.**
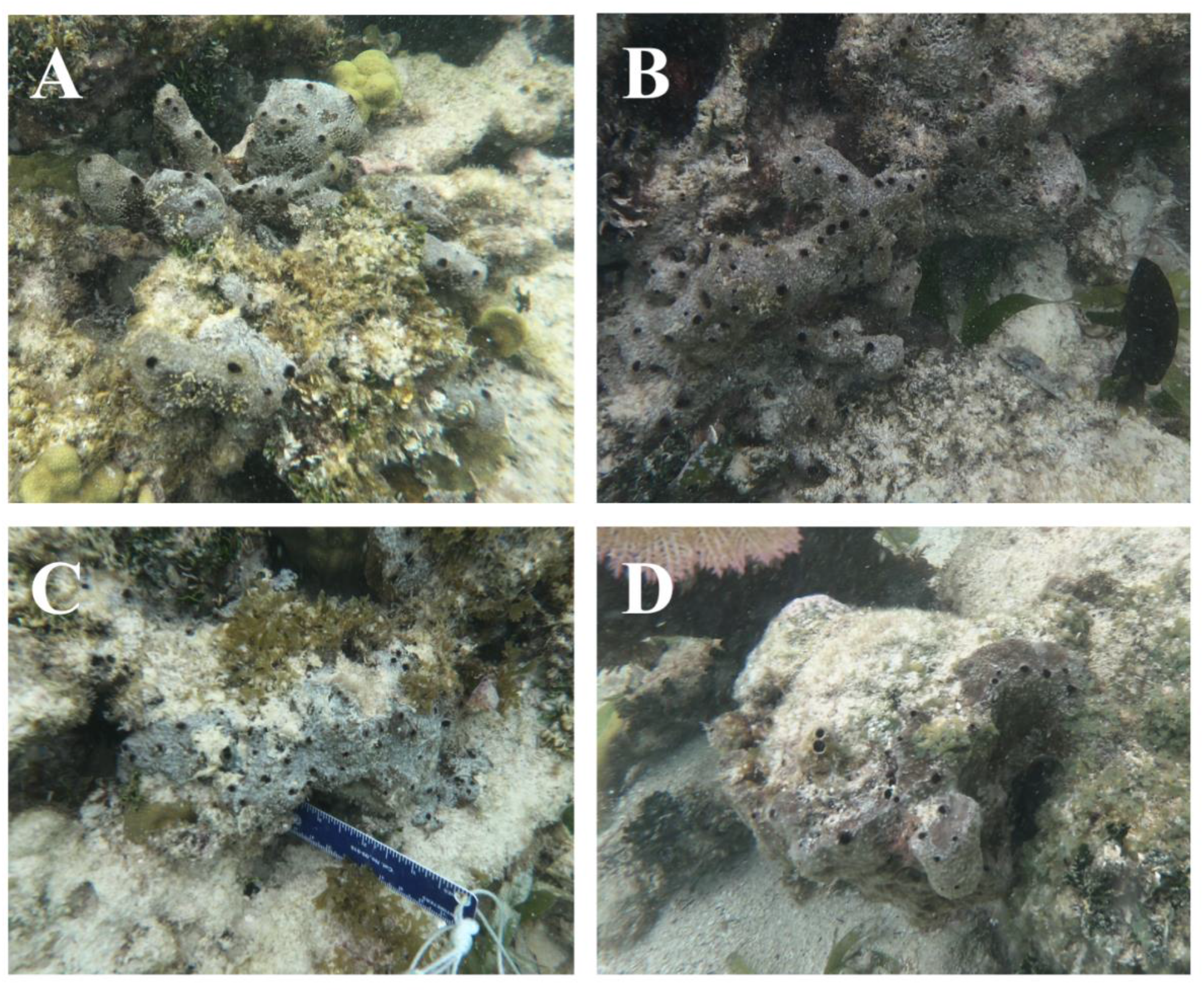
*Ircinia rutzleri* sp. nov. **A:** JK18×23 (holotype), **B:** JK18×26 (paratype), **C:** JK18×22 (paratype), **D:** JK18×25 (paratype).

### External morphology

Conules 1-1.3 mm in height. Oscula flush or slightly raised, always black, 0.7-1 cm in diameter.

### Interior morphology

Massive fascicular fibers 120-240 μm wide, heavily cored and tightly bound. Interconnecting fibers 30-50 um wide, lightly cored. Irciniid filaments 7.5-9 μm wide. Cortex contains abundant inclusions of sand grains.

### Ecology

This species is found on patch reefs co-inhabited by *I. strobilina* in shallow depths (0.5-1 m) adjacent to mangrove hammocks inhabited by sp1.

### Etymology

This species is named for the sponge researcher Klaus Rützler.

### Remarks

Referred to as the ‘Sp. 2’ growth form in Kelly *et al*. (2020).

## *Ircinia* cf. *reteplana* Topsent, 1923

**Representative specimens: JK18×6 (**USNM ######## pending), **JK18×7 (**USNM ######## pending), **JK18×9 (**USNM ######## pending), **JK18×10 (**USNM ######## pending), **JK18×14 (**USNM ######## pending)

**Collection locality: Summerland Key, Florida**

**Note: lodging of museum vouchers pending.**

### Diagnosis

*Ircinia* with a flattened, branching morphology. Branches are usually not interconnecting (Figure 8).

**Figure 8.**
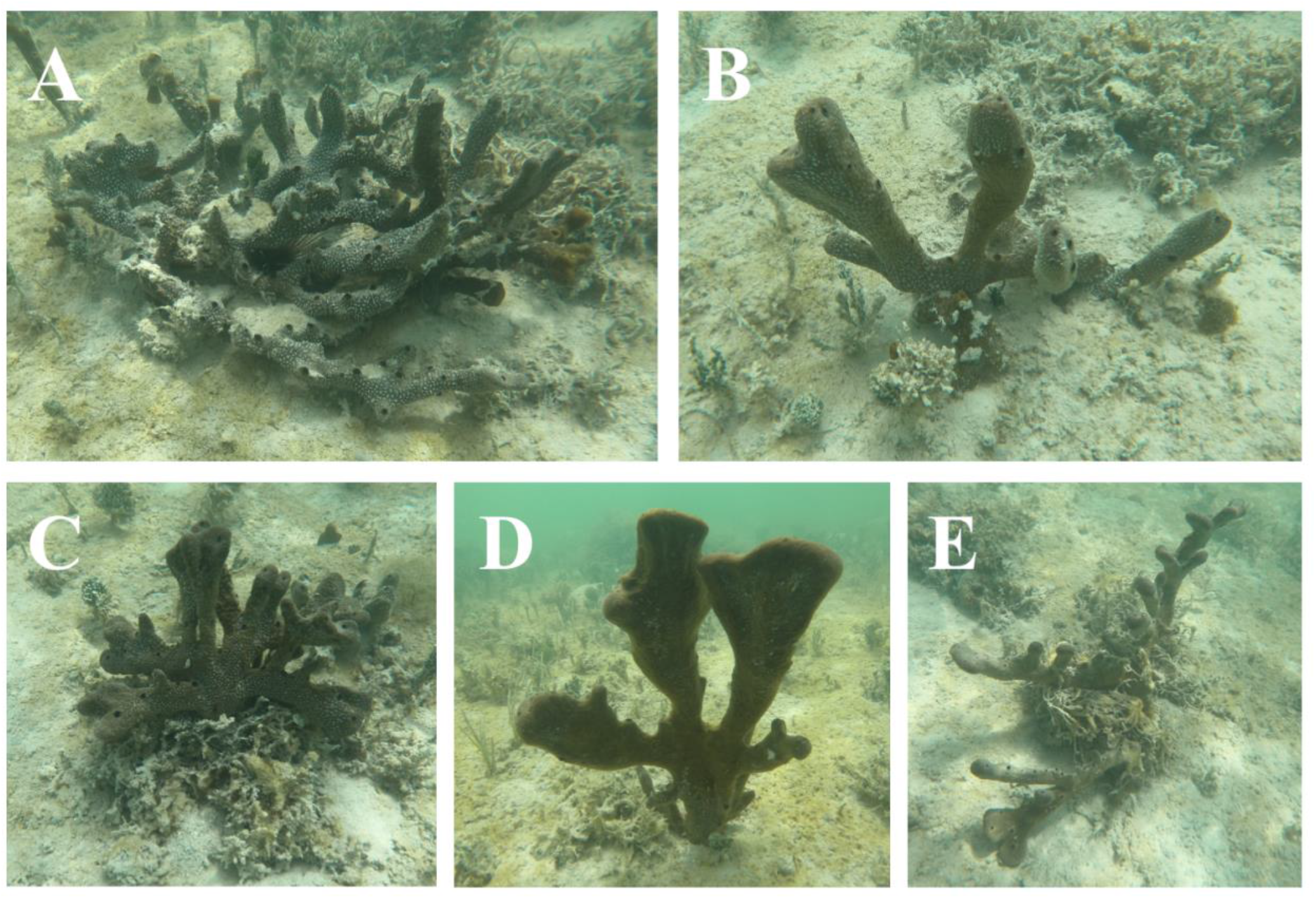
*Ircinia* cf. *reteplana* Topsent, 1923. **A:** JK18×9, **B:** JK18×7, **C:** JK18×6, **D**: JK18×10, **E:** JK18×14.

### External morphology

Surface texture is smooth with 1-1.2 mm-high conules. Most oscula are around 0.5 cm in diameter and are found across the face of the branches, where they sit flush, as well as at the edges of the branches.

### Interior morphology

Massive fascicular fibers are tightly bound, 90-200 μm wide, and heavily cored. Interconnecting fibers 20-80 um wide, lightly cored with spicules and sand. Irciniid filaments 7.5-9 μm wide. Cortex routinely incorporates sand and foreign spicule fragments.

### Ecology

Specimens were collected from a shallow (0.5-1 m) *Thalassia*-dominated seagrass bed and co-occurred next to *I. campana*, sometimes within a meter of each other.

### Remarks

Eukaryotic commensals are mostly crustaceans and polychaetes. Referred to as the ‘Ramose’ growth form in Kelly *et al*. (2020).

### Outlook

Understanding sponge biodiversity is imperative to the protection of tropical marine habitats given the multitude of core ecological functions sponges perform (Bell, 2008; Diaz & Rützler, 2001; Wulff, 2001). *Ircinia* spp. are among the most abundant and ecologically influential sponges on Caribbean reefs although they are also, unfortunately, among the most susceptible to environmental perturbations (Wulff, 2006, 2013). Here, we have demonstrated that disentangling species boundaries within *Ircinia*, arguably one of the most taxonomically challenging sponge genera, can be accomplished with high confidence using an integrative taxonomic framework that evaluates morphology, microbiome composition, and genome-wide SNP data (Kelly & Thacker 2020, *in review*; Kelly et al. 2020, *in review*). The adoption of these data criteria in future species delimitation studies could help further describe species richness within this ecologically important genus and ultimately help science document and defend the hidden biodiversity in sponge fauna.

## Acknowledgements

We thank Dr. Jackie L. Collier and Dr. Liliana Dávalos-Alvarez for their comments on the manuscript; the staffs of the Smithsonian Tropical Research Institute’s Bocas Research Station, the Smithsonian’s Carrie Bow Cay Field Station, and the Mote Marine Laboratory and Aquarium’s Elizabeth Moore International Center for Coral Reef Research & Restoration for their support on logistical aspects of the field work and for helping to organize collection permits. We thank the Ministry of the Environment of Panama; the National Oceanic and Atmospheric Administration’s Office of National Marine Sanctuaries; the Florida Fish and Wildlife Conservation Commission; and the Belize Fisheries Department’s Ministry of Agriculture, Fisheries, Forestry, the Environment & Sustainable Development for granting scientific research permits. We would like to thank Barrett Brooks and Dr. Karen Koltes for help with specimen collections in Carrie Bow Cay. This study was supported by a Fellowship of Graduate Student Travel (Society for Integrative and Comparative Biology) and Dr. David F. Ludwig Memorial Student Travel Scholarship (Association for Environmental Health and Sciences Foundation) awarded to J.B.K. and by grants awarded to R.W.T. from the U.S. National Science Foundation (DEB-1622398, DEB-1623837, OCE-1756249).

## References

Bell, J. J. (2008). The functional roles of marine sponges. Estuarine, Coastal and Shelf Science, 79, 341–353. https://doi.org/10.1016/j.ecss.2008.05.002

de C. Cook, S., & Bergquist, P. R. (1999). New species of dictyoceratid sponges from New Zealand: Genus Ircinia (Porifera: Demospongiae: Dictyoceratida). New Zealand Journal of Marine and Freshwater Research, 33(4), 545–563. https://doi.org/10.1080/00288330.1999.9516899

Degnan, J. H., & Rosenberg, N. A. (2006). Discordance of Species Trees with Their Most Likely Gene Trees. PLOS Genetics, 2(5), e68. https://doi.org/10.1371/journal.pgen.0020068

Diaz, C., & Rützler, K. (2001). Sponges: an essential component of Caribbean coral reefs. Bulletin of Science Marine, 69(2), 535–546.

Diaz, M. (2005). Common Sponges from Shallow Marine Habitats from Bocas del Toro Region, Panama. Caribbean Journal of Science, 41, 465–475.

Erpenbeck, D., Galitz, A., Ekins, M., Cook, S. de C., van Soest, R. W. M., Hooper, J. N. A., & Wörheide, G. (2020). Soft sponges with tricky tree: On the phylogeny of dictyoceratid sponges. Journal of Zoological Systematics and Evolutionary Research, 58(1), 27–40. https://doi.org/10.1111/jzs.12351

Erwin, P. M., & Thacker, R. W. (2007). Incidence and identity of photosynthetic symbionts in Caribbean coral reef sponge assemblages. Journal of the Marine Biological Association of the United Kingdom, 87(6), 1683–1692. https://doi.org/10.1017/S0025315407058213

Hooper, J., & van Soest, R. (2002). Systema Porifera: A guide to the classification of sponges. In Invertebrate Systematics - INVERTEBR SYST (Vol. 18). Kluwer Academic/ Plenum Publishers. https://doi.org/10.1007/978-1-4615-0747-5

Leaché, A. D., Fujita, M. K., Minin, V. N., & Bouckaert, R. R. (2014). Species delimitation using genome-wide SNP Data. Systematic Biology, 63(4), 534–542. https://doi.org/10.1093/sysbio/syu018

Pöppe, J., Sutcliffe, P., Hooper, J. N. A., Wörheide, G., & Erpenbeck, D. (2010). COI barcoding reveals new clades and radiation patterns of Indo-Pacific sponges of the family Irciniidae (Demospongiae: Dictyoceratida). PloS One, 5(4), e9950–e9950. https://doi.org/10.1371/journal.pone.0009950

Riesgo, A., Pérez-Portela, R., Pita, L., Blasco, G., Erwin, P., & Lopez-Legentil, S. (2016). Population structure and connectivity in the Mediterranean sponge Ircinia fasciculata are affected by mass mortalities and hybridization. Heredity, 117(6), 427–439. https://doi.org/10.1038/hdy.2016.41

Rützler, K., Diaz, M., van Soest, R., Zea, S., Smith, K., Alvarez, B., & Wulff, J. (2000). Diversity of sponge fauna in mangrove ponds, Pelican Cays, Belize. Atoll Research Bulletin, 476, 229–248. https://doi.org/10.5479/si.00775630.467.229

van Soest, R. W. M. (1978). Marine sponges from Curaçao and other Caribbean localities Part I. Keratosa. Studies on the Fauna of Curaçao and Other Caribbean Islands, 56(179), 1–94.

Van Soest, R. W. M., Boury-Esnault, N., Hooper, J. N. A., Rützler, K., de Voogd, N. J., Alvarez, B., Hajdu, E., Pisera, A. B., Manconi, R., Schönberg, C., Klautau, M., Kelly, M., Vacelet, J., Dohrmann, M., Díaz, M.-C., Cárdenas, P., Carballo, J. L., Ríos, P., Downey, R., & Morrow, C. C. (2019). World Porifera Database. https://doi.org/10.14284/359

Wulff, J. (1994). Sponge feeding by Caribbean angelfishes, trunkfishes, and filefishes. In In: Van Soest RWM, van Kempen TMG, Braekman JC (eds) Sponges in Time and Space. Biology, Chemistry, Paleontology. Balkema, Rotterdam, pp 265-271. http://www.marinespecies.org/porifera/porifera.php?p=sourcedetails&id=278319

Wulff, J. (2001). Assessing and monitoring coral reef sponges: why and how? Bulletin of Science Marine, 69(2), 831–846.

Wulff, J. (2006). Rapid diversity and abundance decline in a Caribbean coral reef sponge community. Biological Conservation, 127(2), 167–176. https://doi.org/https://doi.org/10.1016/j.biocon.2005.08.007

Wulff, J. (2013). Recovery of Sponges After Extreme Mortality Events: Morphological and Taxonomic Patterns in Regeneration Versus Recruitment. Integrative and Comparative Biology, 53(3), 512–523. https://doi.org/10.1093/icb/ict059

